# Concerns regarding vaccination as a management strategy against koala retrovirus

**DOI:** 10.1101/2021.05.10.443397

**Authors:** Briony A. Joyce, Daniel Watterson, Ariel Isaacs, Keith J. Chappell

**Affiliations:** School of Chemistry and Molecular Biosciences, The University of Queensland, St. Lucia, Queensland, Australia; Australian Institute for Bioengineering and Nanotechnology, The University of Queensland, St. Lucia, Queensland, Australia; Australian Infectious Disease Research Centre, The University of Queensland, St. Lucia, Queensland, Australia

## Abstract

Koalas are of great cultural, ecological and economic significance to Australia. However, they are in steep decline throughout much of their geographic range, with the recently endogenized koala retrovirus (KoRV) presumed to be a contributing factor. Olagoke, et al. ^1^ have proposed vaccination against KoRV as a suitable control mechanism to reduce the severity of KoRV infections and suggest that their recombinant vaccine is effective at invoking a neutralising immune response and significantly reducing viral loads. Here, we report the findings of our own attempts to detect antibodies specific to the KoRV envelope protein (Env) in infected animals. Antibodies against Env were undetectable in all animals, as would be expected in the case of an endogenous virus antigen which should proceed towards a pathway of self-tolerance. This finding calls into question the results of Olagoke, et al. ^1^, and prior work by this team ^2,3^. Following a critical review of this work ^1–3^, we have identified a number of assay inaccuracies and possible over interpretations of the data and conclude that further work to develop and test a vaccine against KoRV in koalas is not supported by evidence.

---

Vaccination against koala retrovirus (KoRV) has been proposed as a control mechanism to reduce disease associated with KoRV infection within koalas. As KoRV is an endogenous virus, integrated within the germline DNA of all koalas throughout the northern states of Queensland and New South Wales, it would be expected that an antibody response to KoRV should proceed towards a pathway of self-tolerance. Despite this, Olagoke *et al*., ^1–3^ have found koalas to have pre-existing levels of anti-KoRV antibodies that can be boosted by vaccination and also that these antibodies capable of virus neutralisation by a KoRV replication assay ^1,3^. It is worth pointing out that the antigen utilised by Olagoke et al. ^1–3^ for ELISA, and as the vaccine, consists of Env amino acids 101 to 659 and is recombinantly expressed as a Glutathione S-Transferase (GST) fusion protein in *E. coli* with no trimerization domain. Of note, Env is expected to exist on the viral surface as a trimer with each monomer predicted to include nine separate disulphide bridges and six potential N-linked glycosylation sites. Due to the lack of enzymes responsible for disulphide bond formation and glycosylation in the prokaryote expression system, along with the absence of a trimerization domain, this protein would be expected to be substantially misfolded and adopt a conformation that poorly resembles that present on the native virus.

To investigate this further, we expressed and purified recombinant Env stabilised with a molecular clamp trimerization domain previously shown to stabilise the native, prefusion structure of class I fusion proteins ^4–6^. This protein was expressed in a mammalian expression system (ExpiCHO; ThermoFisher), enabling correct formation of internal disulphide bonds and native glycosylation patterns ^4^. *In vitro* analyses of the clamp-stabilised KoRV envelope (Env-clamp), including size-exclusion chromatography, electron microscopy and SDS-PAGE, suggested that the Env-clamp antigen was trimeric and adopted a conformation consistent with the expected native KoRV Env pre-fusion structure (Figure 1). Positive control sera against KoRV Env was then generated following Env-clamp immunisation of a rabbit and mice. This stabilised Env-clamp protein was utilised to determine the levels of naturally occurring anti-Env antibody levels within both wild and captive south-east Queensland koalas (*n* = 8) via ELISA and western blot (WB; Figure 2). No anti-Env antibodies were detected within sera from any of the koalas out to a limit of detection (LoD) of 1 in 100 (Figure 2a). Assay controls included quantification of total koala IgG and the immune response elicited to Env-clamp in both mice and rabbits.

**Figure 1.**
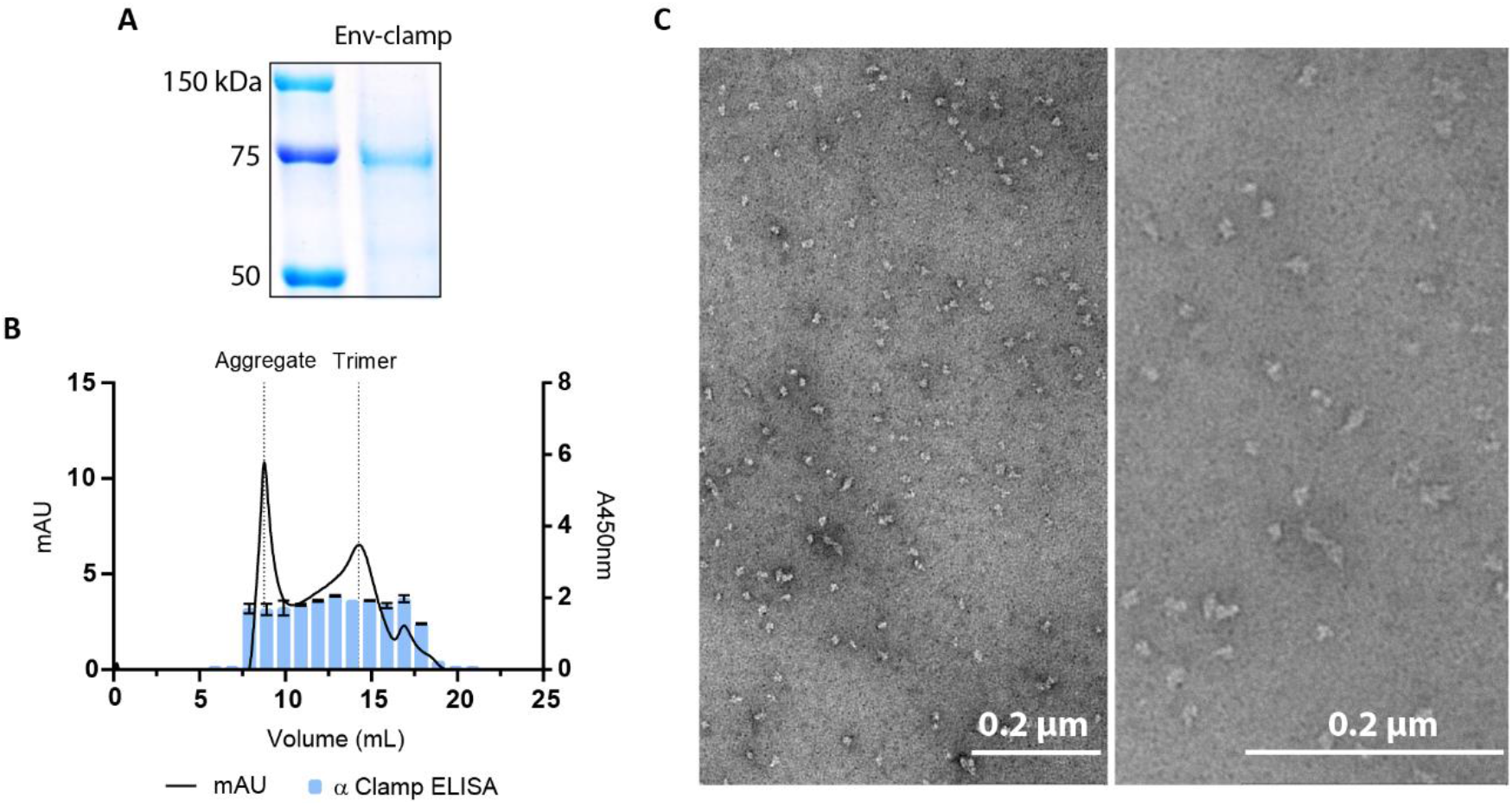
Env-clamp structural characterisation. **A.** SDS-PAGE to assess antigen purity. KoRV-A Env-clamp proteins were expressed in ExpiCHOs, purified via immunoaffinity and run on an SDS-PAGE (4-12%). Precision Plus All Blue Protein Standard (Bio-Rad) used as marker. **B.** Size-exclusion chromatography and ELISA to determine antigen oligomeric state. KoRV Env-clamp proteins were analysed on a Superdex 200 increase 10/300 GL column (GE Healthcare) at an absorbance of 280nm. Fractions were analysed in an ELISA with anti-clamp antibodies. **C.** TEM imaging of trimeric Env-clamp proteins. Proteins from appropriate SEC fractions were negatively stained with uranyl acetate and imaged on a Hitachi HT7700 electron microscope at ×100k (left) and ×200k (right) magnification.

**Figure 2.**
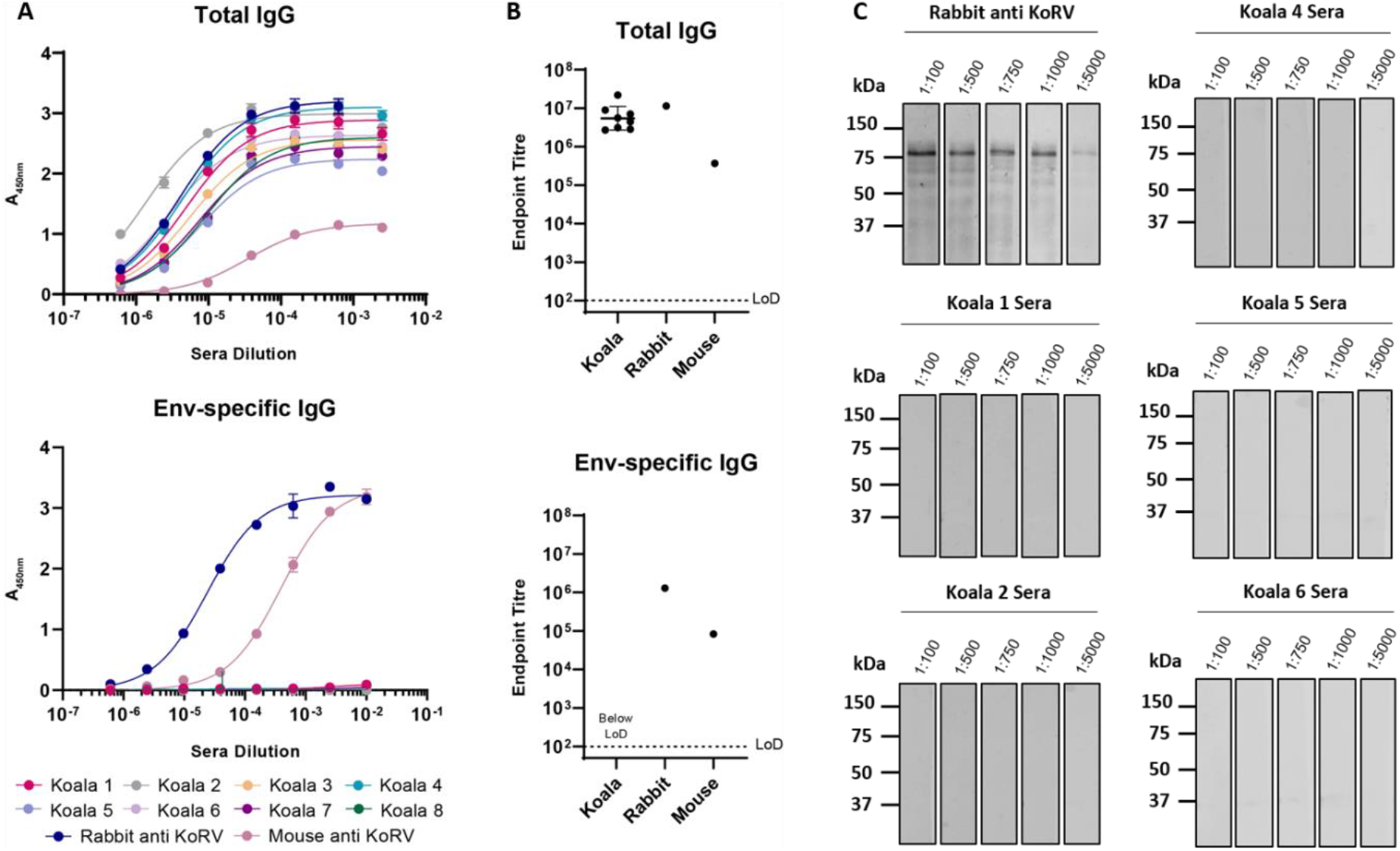
Natural anti-KoRV IgG levels circulating in koala sera. **A.** Total (top) and KoRV env-specific (bottom) IgG levels detected in koala sera (*n* = 8). Rabbit and mouse anti-KoRV polyclonal serums were included as controls. Data is shown as the mean of duplicate values ± S.D, following the subtraction of sera probed against PBS. **B.** Total (top) vs env-specific (bottom) IgG expressed as endpoint titres. Data represents the mean (where applicable) ±S.D. The limit of detection (LoD) is denoted by a dotted line. Endpoint set arbitrarily at 0.1. **C.** Western blots of reduced and denatured KoRV env-clamp probed with titrated koala sera. Rabbit anti-KoRV sera was included as a positive control.

For comparison, Olagoke, et al. detected anti-Env in unvaccinated/pre-vaccinated animals using *E. coli* expressed Env-GST at endpoints of approximately 5,000 ^1^, 1,000 - 8,000 ^2^, and 100 - 2,500 ^3^. Therefore, in all three publications, antibody levels are consistently higher than the LoD in our assay. In Olagoke, et al. ^2^, the assay background is ascribed based on non-infected koala sera and a value of ~1,200 is defined. The fact that the average response in infected koalas is only 2-3 fold the assay background for uninfected animals, raises the possibility that the authors may be over interpreting higher background readings and calls into question the validity of this assay methodology. This is further exemplified in the contradictory results of Olagoke, et al. ^3^ which suggests that KoRV negative animals possess comparable anti-KoRV IgG levels to KoRV positive koalas, as well as a prior publication by some members of the same team, in which no anti-KoRV immune response was identified ^7^.

In each of two vaccination studies, Olagoke, et al. ^1,3^ administered two sequential doses four weeks apart of 50μg of *E. coli* expressed Env-GST formulated with Tri-adjuvant consisting of 250μg polyphosphazine, 500μg host defence peptide 1002, and 250μg poly I:C (Vaccine and Infectious Disease Organization, Saskatchewan, Canada). Following vaccination, a modest 16-fold increase in antigen-specific response was observed ^1^. Notably, large immune responses (≥10^6^ fold) have been previously generated in koalas against a Chlamydia subunit vaccine ^8^, using a highly similar adjuvant formulation. This relatively weak anti-KoRV immune response seen by the authors therefore suggests that koalas are tolerant to the antigen, identifying it as ‘self’, due to the endogenous status of KoRV. Olagoke, et al. ^1,3^ do not determine whether the response is elicited against the KoRV Env or GST portions of the chimeric antigen.

As well as the recombinant *E. coli* expressed Env protein ELISA, Olagoke, et al. ^1–3^ also use a peptide specific ELISA. The authors report various peptide displayed fluorescence intensities above background, however it is not clear from the reported methods exactly how background was calculated. Non-specific binding of monoclonal antibodies to peptides differs depending on amino acid content and so care must be taken with defining background in such assays. This again leads us to speculate that the authors may again be over-interpreting assay background readings. To assess whether koalas may be producing an immune response to linear epitopes within KoRV Env, we performed a WB that was revealed with either serum from a rabbit that had been vaccinated with KoRV Env-clamp or serum from five different koalas. While the rabbit sera could detect the band corresponding to KoRV Env out to a dilution of 1:5000, none of the five koala serum samples tested detect KoRV Env, even at the highest concentration tested of 1:100 (Figure 2c).

Olagoke, et al. ^1,3^ additionally propose that antibodies elicited by vaccination are capable of virus neutralisation, however the assay methodology utilised as the basis of this claim also raises some concerns. Neutralisation titres are reported as the serum dilution that resulted in a 50% reduction in KoRV-A proviral integration as measured by quantitative PCR (qPCR) 65hrs after infection ^1^. A 50% reduction in integration equates to an increase in Ct value of one, which is extremely difficult to accurately assess. Of most concern, Olagoke, et al. ^1,3^ do not report technical nor biological replicate variability and do not define the accuracy of this neutralisation assay.

It is also important to note that the assay methodology and study design both lack appropriate controls. Virus neutralisation was assessed relative to a no serum control, which does not account for any non-specific background neutralisation levels which may present in naïve serum ^1^. Furthermore, the study design did not include a placebo or non-vaccinated control group of animals, which would account for any environmental or seasonal variation ^1^.

Overall, the findings from the neutralisation assay are potentially misleading. Neutralisation titres reported as absolute values without assessment of standard error, mean neutralisation titres reported within the group which may be skewed by high value outliers, and the incorrect use of a statistical method (Student’s T-test) makes the findings appear more significant than in actuality ^1^.

In promoting the therapeutic potential of their vaccine, Olagoke, et al. ^1^ state that pre-vaccination KoRV-A viral loads were completely cleared following vaccination. Here again values are presented as absolutes without acknowledgement of standard error. Additionally, this data is presented as a percentage of the pre-vaccination levels, with values falling below an undefined lower limit of quantification presented as zero. Presentation of data this way is, again, potentially misleading, as the magnitude of the decrease in viral load from pre- to post-vaccination is not able to be quantified and a reliable assessment of statistical significance cannot be determined.

It is well accepted that due to endogenous retroviruses being carried through the germline into each and every somatic cell, they are considered “self” and adaptive immune cells (T and B cells) are rendered tolerant to these retroviral antigens during their development so as to avoid autoimmune disease ^9^. Our findings suggesting the complete absence of any substantial immune response against KoRV Env are consistent with immune tolerance of the endogenous virus. We therefore suggest that vaccinating koalas against the KoRV envelope protein has the potential to cause a break in tolerance and induce auto-immunity, as is proposed for other related retroviruses ^10^. Whilst such a response may not arise following vaccination against the exogenous KoRV subtypes, this is unlikely given these antigens only differ by ~40 amino acids in the receptor binding domain.

Based on our own analysis and detailed assessment of the experimental methods and result reported by Olagoke, et al. ^1–3^, we believe there is little compelling evidence to support vaccination as an approach to protect koalas against disease associated with KoRV. Furthermore, vaccination with an endogenous retrovirus antigen has the potential to trigger autoimmunity and associated pathology, which may be harmful to vaccinated animals. In light of this, we suggest that the findings of Olagoke, et al. ^1,3^ are re-evaluated before any further vaccination studies are initiated in koalas.

## Author contributions

Conceptualisation, K.J.C and D.W. Experimental, B.A.J, D.W and A.I. Article review and editing, B.A.J., A.I and K.J.C.

